# Supervised and unsupervised machine learning methods for modelling current and future habitat of Peruvian anchovy

**DOI:** 10.1101/2025.05.08.652876

**Authors:** Mariana Hill, Tianfei Xue, Jaard Hauschildt, Mariano Gutiérrez, Tronje Kemena

## Abstract

Understanding the drivers and potential impacts of environmental variability on the distribution of Peruvian anchovies, the largest single-species fishery on the planet, is essential for their proper management in a changing world. However, the intricate interactions of these organisms and environmental variability require the use of complex models such as machine learning methods. In this study, we compared three methods for producing habitat maps of anchovies: the traditional Generalised Additive Models, the XGBoost which is a form of supervised machine learning and a new method based on clustering water types as a form of unsupervised machine learning. We optimised the three methods with a parameter grid search algorithm and compared their capability to replicate the mean state of anchovies by comparing them with presence-absence observations along the Peruvian coastline between 1990 and 2010. We used the output of a physical-biogeochemical model as input for the habitat models to produce distribution maps of anchovy. All models successfully simulated the distribution of anchovies along the Peruvian coastline in normal years and a reduced area of distribution with most of the anchovies in the southern part of the domain during the canonical El Niño 97/98. We then applied the models to predict potential changes in the distribution of anchovies under projected temperature and wind conditions by the end of the century. We observed a reduction in the probability of anchovy occurrence under conditions of higher temperature and weaker winds. Two of the three habitat models predicted a severe maximum decline by 90% (GAM) and 75% (XGBoost) whereby the clustering model predicted a moderate maximum decline in anchovy occurrence by 20%.

## 1 Introduction

Understanding the drivers and potential impacts of environmental variability on the Peruvian anchovy (*Engraulis ringens*) is crucial to assess the potential climate change effects on this fishery of global importance as well as the economic consequences. The Peruvian anchovy is the largest single-species fishery in the world (Chavez et al., 2003; Chavez et al., 2008), accounting for 24 (23 for fish oil) to 47 % of the global fishmeal and fish oil supply (Freón et al., 2017; Avadí, Freón, and Tam, 2014) which are an important feed source for aquacultural production (Freón et al., 2014). This species inhabits the coastal waters within the Northern Humboldt Current System (NHCS), typically spanning latitudes from about 4^°^S up to 24^°^S in two well-defined stocks (Bouchon et al., 2021). Anchovies respond in complex ways to the strong environmental interannual variability of the southeastern Pacific Ocean. The anchovy fishery was a proliferating business that increased over the 1960’s reaching landings of 10.9 Mt in 1970 and collapsing shortly afterwards during the strong El Niño event 72/73 (Alheit and Niquen, 2004). Fish populations declined by over 50% during El Niño 82/83 event (Barber and Chavez, 1983) and catches dropped to only 1 Mt in 1998 (Ñiquen and Bouchon, 2004), possibly due to a re-distribution of the fish towards the more coastal waters due to El Niño 97/98 (Bertrand et al., 2004).

Long-term trends in environmental factors due to climate change might substantially impact the habitat of anchovies. Under climate change, the upwelling centre is projected to shift poleward (Rykaczewski et al., 2015), potentially resulting in a displacement of the anchovy population (Pinsky et al., 2013; Checkley, Asch, and Rykaczewski, 2017). However, any wind changes in the NHCS under climate change and their effects on upwelling strength and coastal sea-surface temperature are still highly uncertain, as the effects of increased stratification due to surface warming and increased upwelling-favourable winds may balance each other to some extent (Gutiérrez et al., 2011; Echevin et al., 2012; Echevin et al., 2020). Furthermore, climate change is expected to increase the frequency and intensity of El Niño events (Gergis and Fowler, 2009; Shin et al., 2022) which have affected the fishery in the past.

The drivers of anchovy distribution and its response to environmental changes are not yet completely understood. Chavez et al. (2003) concluded that anchovy abundance may be correlated with cold water regimes, while Bertrand et al. (2011) point out that oxygen also plays an important role. Brochier et al. (2013) and Flores-Valiente et al. (2023) suggest that early stages of anchovy may benefit from climate change as a result of increased retention and potentially higher food availability due to a strengthened stratification, nonetheless, Checkley, Asch, and Rykaczewski, 2017 note that a future reduction in nutrient supply could diminish plankton productivity and alter its composition, thus, as a source of food for anchovies, impacting their population. Currently, anchovies have a competitive advantage compared to other species due to their ability to cope with low oxygen concentrations and have, therefore, been thriving in decades characterised by such conditions (Bertrand et al., 2011). However, under future environmental conditions with even lower oxygen anchovies might be replaced by smaller fish (Salvatteci et al., 2022). This indicates a complex interaction between anchovy and environmental dynamics in the NHCS.

Habitat models, also known as species distribution models or habitat niche models, predict potential areas of distribution of marine organisms, such as anchovies. By finding relationships between their observed occurrence and environmental drivers, these models offer valuable insight into potential climate change-driven habitat changes. To do so, habitat models can be trained using several environmental predictors such as temperature, bathymetry and wave energy, among others (e.g., Luján Paredes, 2016; Oliveros-Ramos, 2014; Gogina et al., 2016; Schubert, Hukriede, and Karez, 2015; Sequeira et al., 2014) to predict biomass, abundance, presence only and presence-absence data (Guisan and Zimmermann, 2000; Oliveros-Ramos and Shin, 2023; Sequeira et al., 2014; Grüss, Drexler, and Ainsworth, 2014). The relationship between these various predictors and the target variables can be very complex and non-linear, requiring machine learning or other advanced modelling techniques to be captured. Generalised Linear Models (GLM) and Generalised Additive Models (GAM) are commonly used methods for modelling habitat due to their capability of fitting non-normally distributed data and capturing the joint impact of several predictors (Guisan and Zimmermann, 2000). In recent years supervised machine learning algorithms such as random forests, and particularly XGBoost, which is a scalable machine learning system for tree boosting (Chen and Guestrin, 2016), have become a popular method with the rise of artificial intelligence applications to scientific problems, as an alternative to the traditional statistical models. Clustering is a method of unsupervised machine learning in which patterns in the data are identified. Unsupervised learning is commonly used for exploratory data analysis but, to our knowledge, has not been applied for making fish habitat predictions so far. Machine learning methods are suitable to disentangle the complex and non-linear links between environmental parameters and to predict the distribution of marine organisms; hence they might be able to capture the intricate relationship between anchovy habitat preferences and its drivers.

In this study, we compared three methods for predicting the distribution of Peruvian anchovy in the northern Humboldt Current System. We applied XGBoost and GAM, both supervised learning methods and a new approach with unsupervised learning that involves clustering. *In-situ* observations were used to train the models. For climate change estimates, we used a physical-biogeochemical model to predict the anchovy habitat. The model parameter validations were done with model data, hence accounting for biases that might arise from the differences between the physical-biogeochemical model output and field observations. We designed a common algorithm for training and comparing the three methods. We compared the predictions by our three types of models during normal years between 1990 and 2010 and 1998 where anchovy catches were low due to El Niño (Bertrand et al., 2004). Finally, we used our models to simulate potential anchovy distribution under idealised climate change conditions.

## 2 Methods

### 2.1 Data description

The observational data was obtained from acoustic surveys performed by the Instituto del Mar del Perú (IMARPE) along the Peruvian coastline in transects up to 200 nautical miles (nm) long from the coast towards the open ocean (Figure 1) from 1990 to 2010 (Gutiérrez et al., 2012). The cleaned dataset consists of 219 thousand samples. Anchovy abundance data is highly skewed to the absence class with values as high as 182.5 thousand but a mean of only 216 m^2^/square nautical mile (nm^2^). Therefore, we transformed anchovy biomass to presence-absence which resulted in 25.3% of the samples with anchovy presence and 74.7% with absence (Figure 1). In addition, we transformed all environmental variables to a range from 0 to 1 using the *StandardScaler* function of *scikit-learn* (Pedregosa et al., 2011). For the supervised learning models, we transformed the distributions of the environmental variables with a quantile transformer into a Gaussian-like distribution (Pedregosa et al., 2011).

**Figure 1.**
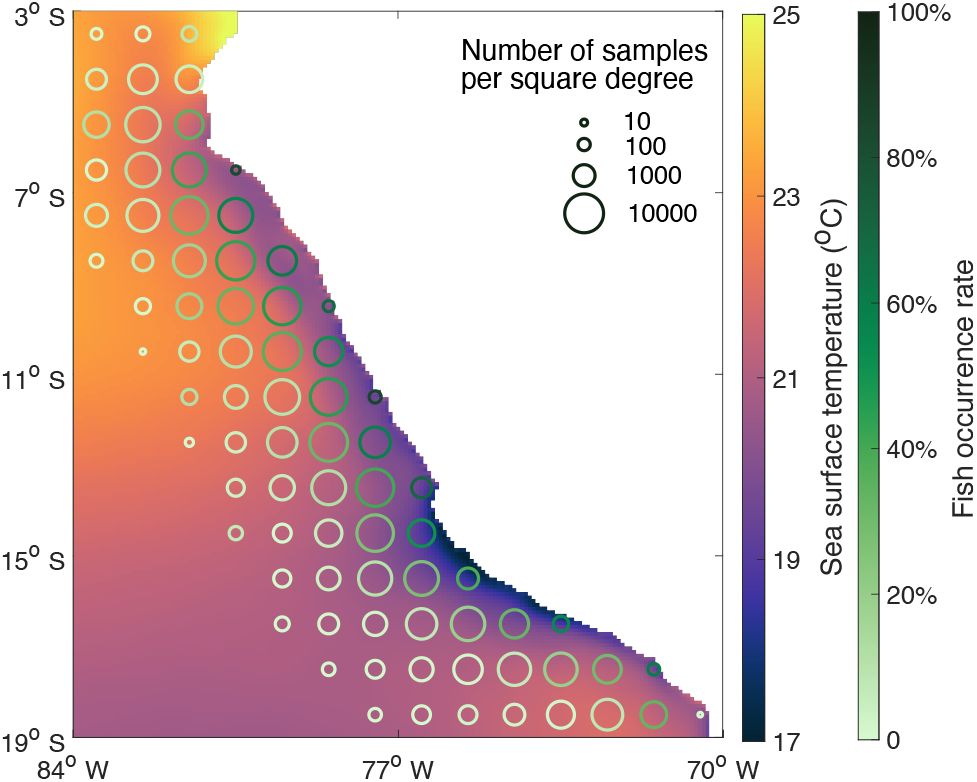
Map of the study area with averaged modelled sea surface temperature (background colour, ^°^*C*) in CROCO-BioEBUS and circles on top showing the occurrence rate of anchovies in acoustic surveys (circle colour) and the total number of measurements available for the period of interest across the sampled region (circle size).

### 2.2 Ocean Physical-Biogeochemical Model

We employed a coupled ocean physical-biogeochemical model to investigate how the environmental conditions affect the distribution of anchovy. The model combines the CROCO (Coastal and Regional Ocean COmmunity model, Shchepetkin and McWilliams, 2005) for physical environmental conditions with the BioEBUS (Biogeochemical model for the Eastern Boundary Upwelling Systems, Gutknecht et al., 2013) model. The CROCO framework provides the background for simulating ocean circulation, temperature, and salinity on a regional scale that extends from 10^°^N to 33^°^S and from 69^°^W to 118^°^W with a horizontal resolution of 1/12 ^°^ and 32 vertical sigma levels. The BioEBUS model is integrated with CROCO to represent the biogeochemical environment with a complex nitrogen cycle including oxygen-dependant processes like nitrification and remineralisation. The physical component of the model runs in time-steps of 500 seconds, the biogeochemical component in steps of 1500 seconds and the model output has a monthly resolution. Here, to align with the acoustic cruise data detailed in Sect. 2.1, we have narrowed the focus of our study to a smaller domain covered by the available observations, ranging from 3^°^S to 19^°^S latitude and from 70^°^W to 84^°^W longitude, as illustrated in Fig. 1.

#### 2.2.1 Hindcast simulation

To simulate the historical interannual variability in the environment that potentially affects fish distribution, we used an interannual configuration of CROCO-BioEBUS. Surface heat and freshwater flux forcing for CROCO are provided by CFSR (Climate Forecast System Reanalysis) data, and the wind forcing by the CCMP (Cross-Calibrated Multi-Platform) product with a temporal resolution of six hours. Initial and boundary conditions are from monthly SODA (Simple Ocean Data Assimilation, Carton, Chepurin, and Chen, 2018) and monthly climatology CARS (CSIRO - Commonwealth Scientific and Industrial Research Organisation Atlas of Regional Seas, Ridgway, Dunn, and Wilkin, 2002). A 30-year spin-up period, by repeating the forcing of the year 1990, ensures model equilibrium before conducting simulations for the period from 1990 to 2010. After spin-up, the model is forced by monthly interannually-varying forcing during the period from 1990 to 2010, which was later used for training the habitat models. The same model set-up was previously used in an End-to-End model study investigating the bottom-up impact of plankton dynamics on higher trophic levels, including anchovy (Hill Cruz et al., 2022). Model evaluations are available in (Hill Cruz et al., 2022; Xue et al., 2024 [In review]).

We sampled the model output at the same spatial and temporal coordinates where field observations were collected. Sea surface temperature (SST), distance from the coast and depth are well correlated between the model and observations (Table 1; Figure 2). Meanwhile, sea surface salinity and oxycline depth have a lower correlation of 0.63 and 0.56, respectively. Furthermore, chlorophyll *a* has higher values in CROCO-BioEBUS than in the observations (Appendix A) and it exhibits a correlation of only 0.28 so we did not include this variable in our habitat models. A bi-modal distribution is observed in SST (Figure 2) due to the sampling taking place in two main seasons (see Appendix A). On the other hand, the two peaks are less distinct in the CROCO-BioEBUS model SST, possibly due to the higher diffusion compared to the real world.

**Table 1.** Correlations between environmental variables in observations and in CROCO-BioEBUS.

| Variable | Correlation (r) |
| --- | --- |
| Water depth* | 0.99 |
| Distance from the shelf | 0.96 |
| Distance from the coast* | 0.93 |
| Sea surface temperature* | 0.87 |
| Sea surface salinity | 0.63 |
| Depth of the oxycline* | 0.56 |
| Chlorophyll *a* | 0.28 |

\* indicates variables later used in machine learning methods.

**Figure 2.**
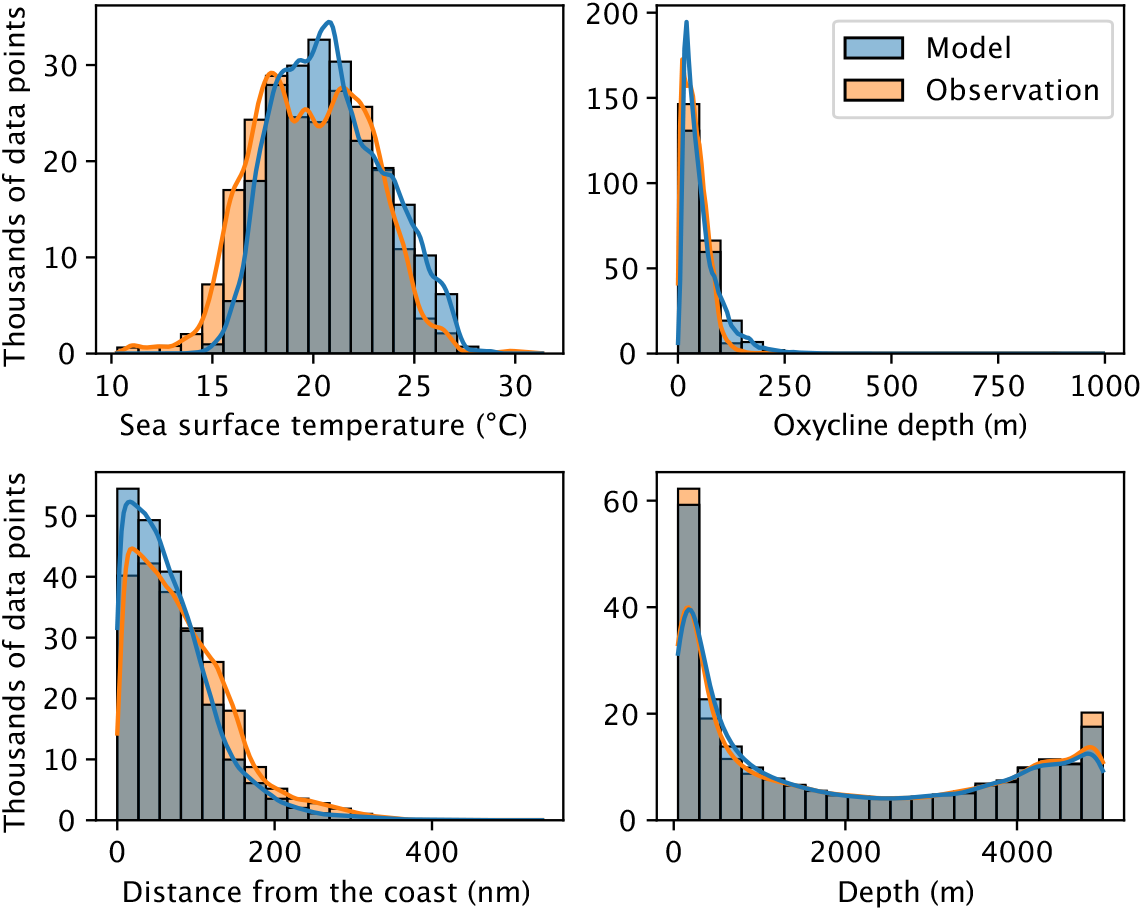
Distribution for observational data and CROCO-BioEBUS hindcast output. The latter is interpolated in time and space to match observations. Lines indicate a kernel density estimate.

**Figure 3.**
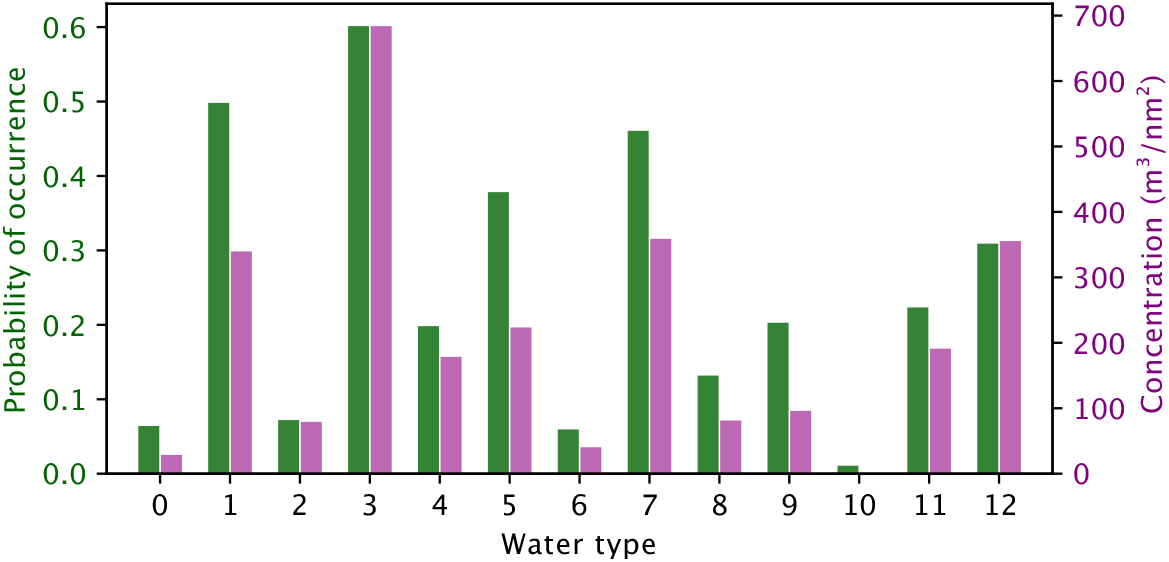
Anchovy probability of occurrence and concentration predicted for each water type.

#### 2.2.2 Climate change experiments

We performed a set of three sensitivity experiments (TEMP+, WIND+, WIND-) to test how our predicted anchovy distribution responds to different environmental states simulated by the coupled physical-biogeochemical model. The sensitivity experiments were designed to represent wind speed changes and warming which could realistically occur by the end of the century according to climate models. These changes were applied to a climatological reference simulation (REFERENCE), which was set up by averaging the forcing of the hindcast simulation described above between 1990 and 2010. The reference simulation was run for a 20-year spin-up period, after which the sensitivity experiments were started. The reference simulation and the sensitivity experiments were then run for a period of 10 years and the last year was used for analysis. Since the climatological forcing contains no interannual variability, the model already reaches a quasi-steady-state within this time-frame in the upper 200-300 m of the ocean, which are relevant for the anchovy habitat.

The warming experiment (TEMP+) was designed to reflect an increase in surface air temperature and water column stratification consistent with end-of-century projections from climate models (Echevin et al., 2020), while keeping the wind forcing constant. We applied a spatially uniform ocean temperature offset at the open boundaries and to the initial temperature fields, in the form of a vertical profile. To compute this average profile of ocean warming, we took the temperature difference between 2006-2100 in the CMIP5 (Coupled Model Intercomparison Project Phase 5, Taylor, Stouffer, and Meehl, 2012) multi-model ensemble average under the RCP8.5 emission scenario and averaged it spatially over our ocean model domain. We then applied the surface value of the previously computed water temperature change profile (2.2 ^°^C) as a spatially uniform offset to the surface air temperature, which is used by the model for computing air-sea heat and momentum fluxes. A similar approach of modifying surface air temperature using the bulk forcing option in CROCO has been used before in regional climate change downscaling simulations to add a forcing perturbation (e.g., Espinoza-Morriberon et al., 2017; Echevin et al., 2020).

To further investigate the uncertainty of potential wind changes and their effect on the anchovy habitat in the region, we increased the wind speed by 10% (WIND+) in a first sensitivity simulation and decreased it by 10% (WIND*−*) in a second one. This corresponds to a wind *stress* change of approximately *±* 20%, as the wind speed enters quadratically into the wind stress parameterisation. All surface fluxes (momentum, heat, CO_2_, O_2_) are calculated by a bulk formula and are thus consistent with these wind changes. Note also that the wind speed changes are small compared to some previously published sensitivity studies (e.g., Echevin et al., 2012; Travers-Trolet et al., 2014) and reflect a plausible range as suggested by global climate models and downscaling experiments in the region (see Echevin et al., 2020, and references therein).

### 2.3 Habitat models

We fitted a series of XGBoost classifiers to the presence-absence observations with the Python package *xgboost* (Chen and Guestrin, 2016). To optimise the parameters for XGBoost, we conducted cross-validations together with a brute force grid search for the parameters: learning rate (0.001 to 0.3), depth (2 to 40), and number of estimators (20 to 700). Out of these 200 optimised model configurations, we picked the best model in reference to two metrics: balanced accuracy (in the case of binary classification area under the ROC curve) and Cohen’s Kappa (see Table 2), which are suitable scores for classification problems with imbalanced classes and habitat modelling (Liu, White, and Newell, 2009; Ben-David, 2008; Wardhani et al., 2019).

**Table 2.**
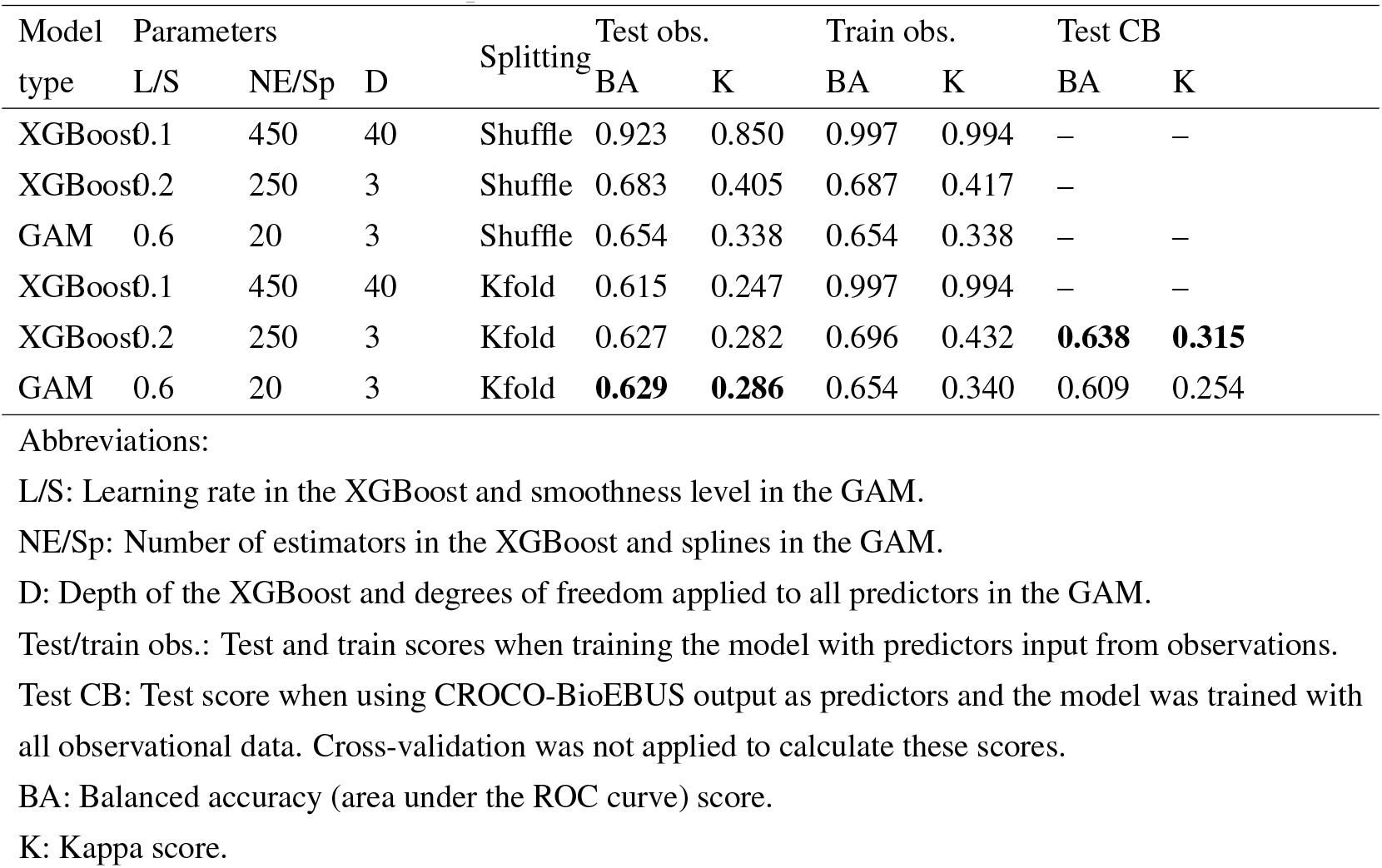
Comparison of XGBoost model and GAM against single observation points. The highest test scores are marked in bold. Shuffle split trained models are not considered for further use (see text).

We compared the performance of these models with Generalised Additive Models (GAM) using the *PyGAM* package (Table 2). We used the *LogisticGAM* model which has a Binomial distribution and a logistic link function with all the default parameters (Servén and Brummitt, 2018). Both XGBoost and GAM were cross-validated with 5 splits applying two types of splitting: standard shuffle and Kfold. We then applied the best models to make predictions of anchovy presence-absence using environmental output from the CROCO-BioEBUS model as predictors.

For the unsupervised learning method, we clustered the data points into water types based on their distance from the coast, SST, bathymetry, depth of the oxycline at 1 ml O_2_ L^−1^ and the month when the sample was taken using Gaussian mixture model probability. Then we calculated the probability of finding anchovy in each water type by averaging the presence (1) and absence (0) points falling within the water type. Similarly, we calculated the average concentration of anchovies in each water type. Calculating total anchovy biomass is beyond the scope of this paper; therefore, we kept the original units from the raw data of m^2^/nm^2^ which is based on acoustic surveys. Since clustering is an unsupervised machine learning method, it is not possible to do cross-validation and due to the applied statistics, we cannot compare to the other models using the standard classification metrics.

In order to compare the three methods, we produced a heatmap of observations averaging at least four data points per grid cell. Then we calculated the mean square root error (RMSE) and correlation coefficient between the grid cells in the observations heatmap and each of our modelled habitat maps. Based on these values, we fine-tuned the parameters of the GAM, XGBoost, and clustering (covariance type full) using brute force grid search. Parameter ranges can be found in Tab. 3. Finally, for the clustering approach, we also optimised a scaling factor applied to the probabilities of finding anchovies in each water type. This had no impact on the correlation but slightly reduced the RMSE. All models took the same environmental predictors as input, as described above.

Once we had decided on the best parameters comparing the models with the hindcast observation heatmaps, we applied each of the three models to simulate a climatology of anchovy probability of occurrence taking as predictors the CROCO-BioEBUS output of the sensitivity studies described in Sect. 2.2.2 to evaluate the climate change impact on their habitat. The XGBoost and clustering methods can only predict habitat when the environmental variables fall within the range that was used to train the models. However, because the interannual variability in the NHCS is high in the hindcast simulation that was used to train the models, the comparatively small changes applied to the climatological simulation to represent climate change fall within the range of the training data.

## 3 Results

### 3.1 Models training

In our initial approach, we performed the parameter grid search for XGBoost with a standard shuffle splitting method, achieving very high train (0.997) and test (0.923) balanced accuracy scores as shown in Tab. 2. Given the potential for spatial or temporal auto-correlation in dense-sampled observational cruises, these results might be overly optimistic (Salazar et al., 2022). When applying the Kfold splitting method to this complex XGBoost model configuration (maximum depth is 40), the test score dropped by over 30%, strongly indicating model overfitting on likely auto-correlated data. To address this, we also conducted a parameter optimisation with a Kfold splitting method, which splits data into k temporally consecutive folds to prevent overfitting from temporal autocorrelation. Among the models with Kfold-split data, the XGBoost configuration with the highest maximum depth (40) yielded the best training scores. However, the considerably lower test scores suggest overfitting. Conversely, a simpler model with a depth of 3 produced closer test and training scores for both Kappa (0.282 and 0.432, respectively) and the balanced accuracy (0.627 and 0.696, respectively)., which were comparable to those of the GAM (Kappa: 0.286, balanced accuracy: 0.340). For further validation, we favoured Kfold splitting, although the GAM model also showed slightly higher test scores when shuffle-splitted, considering this method is more conservative. Following the integration of data from the CROCO-BioEBUS model instead of observations, the balanced accuracy and Kappa for XGBoost slightly improved from 0.627 to 0.638 and 0.282 to 0.315, respectively, while for GAM, they decreased from 0.629 to 0.609 and 0.286 to 0.254, respectively (Table 2). This performance indicates that, despite slight differences, the CROCO-BioEBUS model output is close enough to the observations to produce a reliable habitat map for Peruvian anchovies, demonstrating good generalisation across the two different data sets.

### 3.2 Anchovy response to interannual variability

All three models predict a distribution of anchovies near the coast as seen in the observations. In years other than 1998, the distribution extends over the whole Peruvian coastline. In 1998, the probability of finding anchovy decreases with respect to the average of the other years between 1990 and 2010 and the south of the Peruvian coast remains the main area with high chances of anchovy occurrence (Figure 4). The XGBoost model and GAM show a similar map and the same range of probabilities as the occurrence rate in observations (Figure 4). On the contrary, the clustering map has a narrower range of anchovy probabilities, overestimating in regions where the occurrence rate is low and underestimating in regions where finding anchovies is more likely. However, this method exhibited the highest correlation coefficient (0.527) and the lowest RMSE (0.317, Table 3). The correlation coefficient describes the linear relation of the observation and model data, while the RMSE provides the error. On the contrary, the visual comparison suggests that the XGBoost and GAM are better alternatives than the clustering for predicting anchovy probability of occurrence since they show a wider range of predictions reaching zero offshore and up to 1 near the coast. In contrast, clustering represents better the mean state.

**Table 3.** Comparison of habitat maps produced with XGBoost, GAM and clustering models using predictors input from the CROCO-BioEBUS model. Parameter range for the grid search is indicated in the brackets. The GAM grid search was also conducted on different covariance types: full, tied, diagonal, spherical.

| Model | Parameters | Parameter value | Parameter range | Correlation coeff. | RMSE |
| --- | --- | --- | --- | --- | --- |
| XGBoost | Maximum depth | 2 | 2 - 40 | 0.468 | 0.345 |
|  | Number of estimators | 600 | 20 - 700 |  |  |
|  | Learning rate | 0.03 | 0.001 - 0.3 |  |  |
| GAM | Splines order | 4 | 1 - 5 | 0.384 | 0.360 |
|  | Number of splines | 11 | 6 - 20 |  |  |
|  | Smoothness level | 0.75 | 0.0005 - 0.8 |  |  |
| Clustering | Number of components | 13 | 2 - 30 | 0.527 | 0.317 |
|  | Convergence threshold | 0.0005 | 0.0001 - 0.005 |  |  |

**Figure 4.**
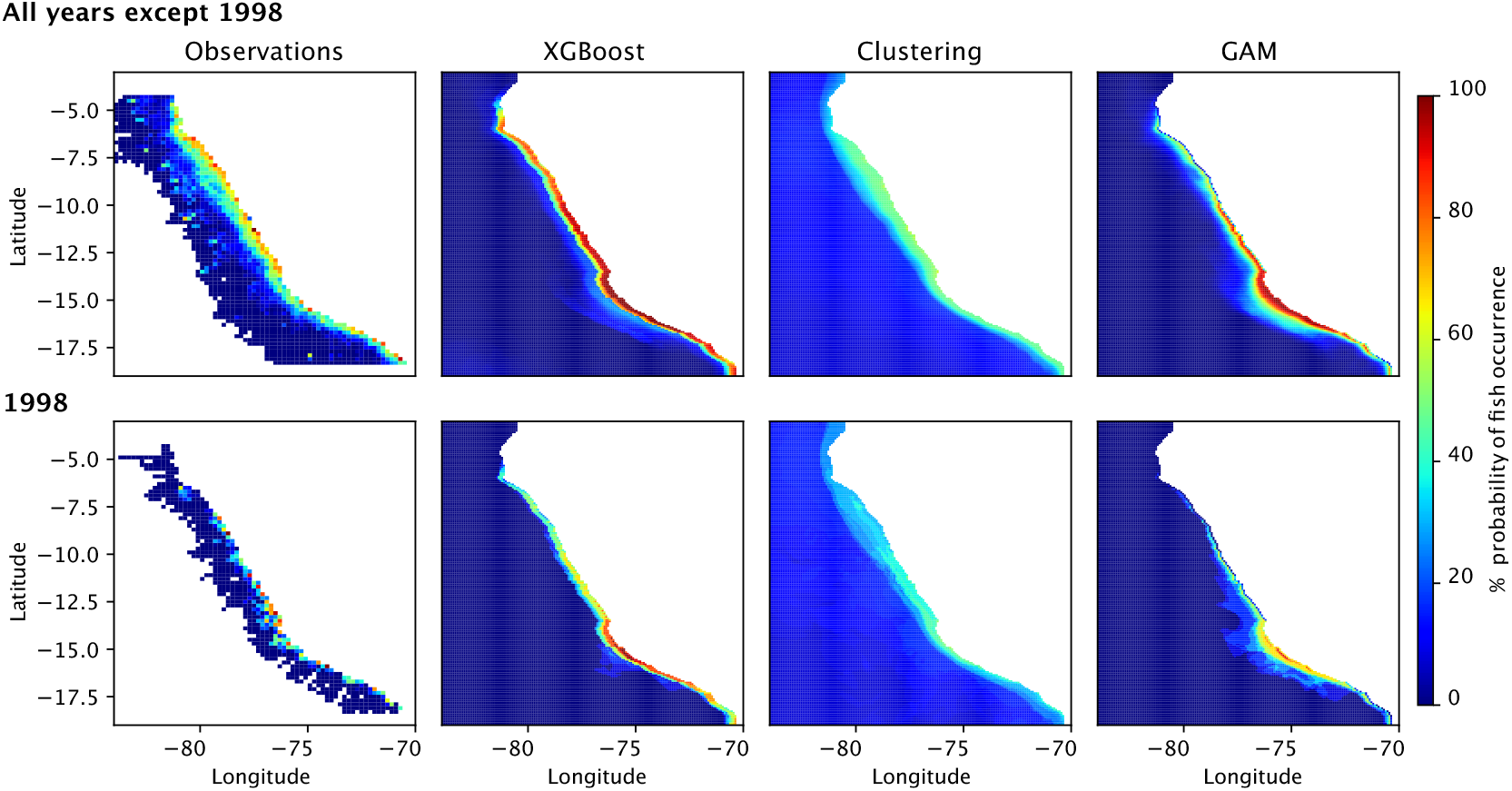
Comparison of anchovy occurrence rate in the observations and probability of occurrence predicted by each habitat model.

The clustering method is also able to predict average anchovy concentration in each water type (Figure 3) while XGBoost and GAM were only trained as classifiers. Although concentration and probability of occurrence generally follow a similar pattern, there are differences in their relative changes across water types (Figure 3).

### 3.3 Anchovy response to climate change

The XGBoost and GAM models predicted a general increase in anchovy probability of occurrence with stronger winds and a decrease in probability with warmer temperatures and weaker winds. A severe maximum decline by 90% (GAM) and 75% (XGB) occurred due to the effect of warmer temperatures. The clustering method predicted similar but weaker responses in the coastal region (Figure 5). Here, a moderate maximum decline in anchovy occurrence by 20% is observed. However, while changes in anchovy occurrence in the open ocean are negligible in the supervised learning models, the relatively high occurrence predicted by the clustering resulted in a high sensitivity to climate change offshore. This is likely due to the very weak overall response of the clustering and relatively high probability of occurrence offshore with respect to the other models which result in a weak signal-to-noise ratio (Figure 5).

**Figure 5.**
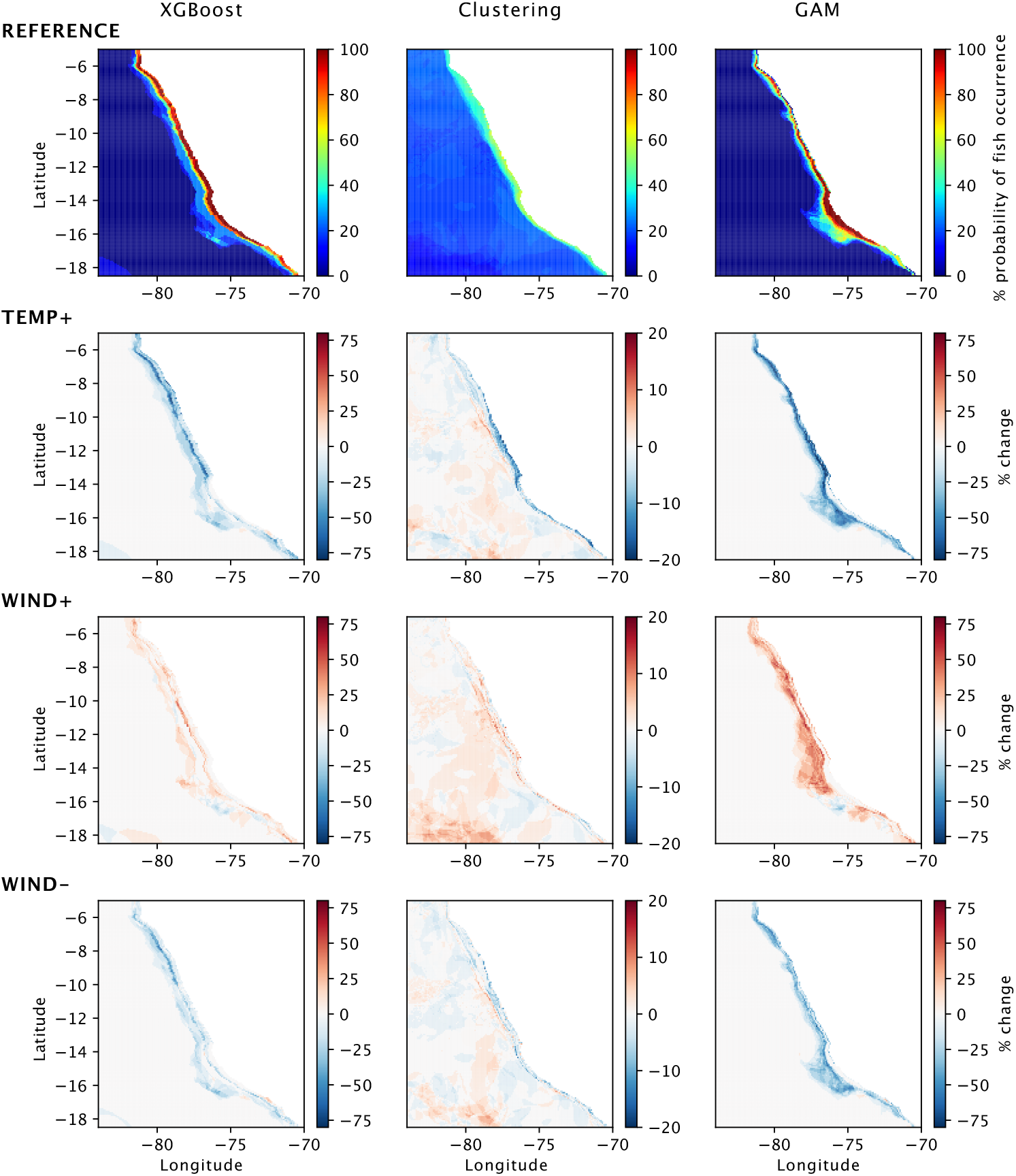
Probability of anchovy presence predicted by each habitat model in the reference simulation (top) and relative change in the climate change scenarios.

## 4 Discussion

The Peruvian anchovy is one of the most studied fisheries due to its economic value but also the complexity of its response to environmental variability. Despite the work done by numerous researchers in the last half of a century, anchovy’s behaviour is still to some extent unpredictable. In this study, we compared three methods for modelling the habitat of Peruvian anchovies. We used a XGBoost model and a GAM which is a statistical method, both trained via supervised learning. These methods have been commonly used to model habitat. We also tested a new method based on defining water types with unsupervised machine learning, or clustering, and associating anchovy occurrence to each of them. We produced habitat maps with all three models using data from a physical-biogeochemical model and compared their capability for producing realistic maps based on two statistical metrics and a visual assessment of the maps. In Sect. 4.1 we discuss the implications of using output from physical-biogeochemical models as predictors for habitat models. In Sect. 4.2 we put the hindcast and climate change predictions from our model in context. Finally in Sect. 4.3 we talk about general challenges for modelling the distribution of marine species and how we addressed them.

### 4.1 Applying a Physical-biogeochemical model as forcing for habitat models

Producing habitat maps depends on the availability of continuous data over the whole region of interest. This can be obtained, for example, from satellites or *in situ* observations. However, satellite data is limited to certain variables that can be detected through remote sensing and *in situ* observations are limited in their spatial coverage. Physical-biogeochemical models, on the other hand, provide a wide range of environmental variables that are not easily observable such as oxycline depth (e.g., Espinoza-Morriberón et al., 2021) and integrated biomass and energy fluxes across the trophic web, which can comprehensively capture the impacts on anchovy feeding and potentially offer a better indicator of fish production (Xue et al., 2024; Friedland et al., 2012). Furthermore, physical-biogeochemical models can be used to conduct sensitivity experiments (e.g., Echevin et al., 2012; Meier et al., 2019) taking into account the changes in all physical and biogeochemical variables that are relevant for the fish habitat such as temperature and oxygen.

We trained and tested the XGBoost model and the GAM with field observations and then coupled them to a regional physical-biogeochemical model to produce habitat maps. We observed similar values in the balanced accuracy and the Cohen’s Kappa score when using predictors from a physical-biogeochemical model in comparison to observational data indicating no loss in predictability due to changing from model data to observations. The XGBoost model and GAM were able to produce very similar maps even when using predictors from a completely different dataset than the one that was used to train them. This indicates that the trained habitat models have high capability to generalise and the relatively low test scores have a different source than the different data domains. Relevant factors could be, for example, the stochastic nature of anchovy appearance (discussed later) or better predictors being needed as model input. However, a similar predictability for both datasets opens the possibility of exploiting the potential of regional physical-biogeochemical models for climate change predictions in the field of habitat modelling.

Earth System Model (ESM) simulations are necessary to predict the impact of climate change on the distribution of marine organisms due to changes in environmental conditions. One way to do this is to add the anomalies simulated by the ESM to the observations used to train the habitat model (e.g., Oliveros-Ramos and Shin, 2023; Sequeira et al., 2014). However, this makes the habitat model susceptible to an unknown error if applied to ESM data since it has been trained and tested only on observations. In addition, the interpolation of global results of the ESM output might not necessarily capture the nuances of a particular region (Echevin et al., 2020). Furthermore, various downscaling and interpolation methods can yield differences in the magnitude of change and in spatial patterns (Pozo Buil et al., 2023). A second method is to use the output of regional models to directly feed habitat models. Nevertheless, the habitat model must be able to generalise well and avoid overfitting to the training data from observations, and be stable enough to handle any noise or bias in the regional model output in order to make good predictions. While regional models also require external forcing from an ESM (e.g., Echevin et al., 2020), some global ESMs such as the Flexible Ocean and Climate Infrastructure (FOCI, Matthes et al., 2020) allow for the implementation of nests with a very fine grid resolution, eliminating the need for downscaling and interpolation at all. Temporal variability in habitat has been suggested to have an impact on how the biomass of marine organisms responds to changes in plankton (Hill Cruz et al., 2022). In addition, fish variability might as well have an impact on the biogeochemistry of the marine ecosystems (Travers-Trolet et al., 2014; Getzlaff and Oschlies, 2017; Hill Cruz et al., 2021; Bianchi et al., 2021). Therefore, being able to run habitat models using direct output from physical-biogeochmeical models opens the possibility for doing dynamic couplings between the two models and capturing the feedbacks between their interaction in end-to-end models.

### 4.2 Climate change and interannual variability impacts on anchovy habitat

Observations indicate that Peruvian anchovies are generally adapted to cold coastal waters with high productivity and are widespread along the entire Peruvian coast Bakun and Broad, 2003; Ñ iquen Carranza et al., 2000. During El Niño, Peruvian anchovies are found to move closer to the coast and migrate southward, rather than maintaining their typical widespread habitat (Mathisen, 1989; Ñiquen and Bouchon, 2004). This aligns with our findings, where only the very coastal region in the southern part of Peru remains a suitable habitat according to the XGBoost model and GAM during 1998 (Fig. 4). The oscillations in the range of the anchovy habitat could potentially influence changes in the fish population; however, as the fish are located further from their typical habitats, they become less accessible for fisheries, leading to an underestimation of fish biomass from commercial catches (Bertrand et al., 2004). Previous studies suggest that environmental changes on different timescales (e.g., interannual and decadal) influence the anchovy habitat in a similar way, which could be indicative of broader climate change impacts (Alheit and Niquen, 2004; Bertrand et al., 2004).

Our climate change experiments indicated a decrease of anchovy occurrence probability along the Peruvian coast with increased temperature and decreased wind. This is in line with Salvatteci et al. (2022) who suggests a replacement of anchovy by smaller fish as a consequence of climate change. In contrast, increased wind energy has the potential to expand the potential habitat for anchovies or counteract the habitat contraction due to warming. Rykaczewski et al., 2015 have suggested that the coastal upwelling zones will be shifted polewards under global warming, resulting in a slightly decreased upwelling strength for the Peruvian coast. However, our simulations did not result in a latitudinal shift of the anchovy habitat for any of the experiments but rather a decrease (TEMP+, WIND-) or increase (WIND+) in the habitat suitability along the entire Peruvian coastline. This is likely a result of our idealised wind speed perturbation, which uses a spatially uniform scaling factor and does not capture potential changes in the wind pattern. Given the uncertainty of projected wind changes in the region, this is a reasonable first assumption.

In this study, the XGBoost model and GAM predicted a similar change in both sign and magnitude in the anchovy distribution as a response to climate change. Such methods can be used for moderate climate change projections based on climatological predictors because, in the NHCS, the strong interannual variability observed in the hindcast under present-day climate conditions is stronger than the climate change impacts reflected in the climatology. However, these methods, as well as the clustering, would not be suitable for predicting extreme changes that fall outside of the range of the training data; for example, in projections where stronger and more frequent El Nino events are considered (Shin et al., 2022), which could cause major changes in anchovy habitat and consequently impact fish population and catches (Arias Schreiber, Ñ iquen, and Bouchon, 2011). Scenarios with a stronger environmental change should rely on simpler methods such as Generalised Linear Models with better capabilities to extrapolate.

Despite the general trend observed in models of a preference for colder waters and the implications for climate change, the habitat of Peruvian anchovies remains somewhat unpredictable since they have shown impressive plasticity in recent years. In 2022, the Instituto Público de Investigación de Acuiculture Pesca (2023) confirmed the presence of anchovies in the Gulf of Guayaquil, in Ecuador, located further north than their usual distribution off Peru. This highlights the adaptable nature of anchovies and the importance of continuous monitoring schemes for ensuring that the most up-to-date changes in species behaviour are captured. Since our habitat model was trained with data until 2010, covering only the Peruvian Exclusive Economic Zone, it might not make reliable predictions for the Gulf of Guayaquil. Strong decreases in temperatures were reported in the Easter Pacific Ocean as well as a shallower thermocline in the Gulf of Guayaquil in 2022 during the La Niña event (Instituto Público de Investigación de Acuiculture Pesca, 2023; Senamhi, 2022), possibly facilitating the occurrence of anchovy further north (Instituto Público de Investigación de Acuiculture Pesca, 2023). Hence, extending the configuration of our CROCO-BioEBUS model and training the habitat models with up-to-date observations is necessary to be able to simulate an updated hindcast of the anchovy habitat in more recent years and have a clearer picture of their potential response and possibilities of adaptation to climate change.

### 4.3 Challenges modelling fish habitat

In order to overcome the challenges of fitting all habitat models used in our study with a common metric and compare both supervised (GAM and XGBoost) and unsupervised (clustering) methods, we opted for producing a heatmap of the observations and comparing the heatmap cells against predicted distribution by the habitat model with input from the physical-biogeochemical model. This method allows us to compare probabilities against occurrence rate rather than single presence-absence points. We were able to train all three models using randomised grid search against the same set of observations and produce maps that provide a realistic distribution for anchovies, concentrating most of them near the coast of Peru and leaving the open ocean free.

A main challenge when modelling marine organisms distributions is the extrapolation of models to regions where observational samples are not available (e.g., Sequeira et al., 2014) or sparse, for instance, the open ocean. This has been handled by adding pseudo-absence data points in areas where it is known that the species of interest does not naturally occur (Luján Paredes, 2016). Another alternative for the prediction of fish occurrence is to use a method that is based on presence-only data, such as maximum entropy models (Bang et al., 2022). However, these models are prone to geographical sampling bias (Syfert, Smith, and Coomes, 2013). Our XGBoost model and GAM were trained without pseudo-absence points showing no extrapolation issues, indicating that these models generalise well on unknown data.

A further obstacle when modelling the distribution of highly mobile marine animals such as fish is that their presence in the sampling does not only depend on the suitability of environmental conditions but also on the chances of sampling at the specific time when the school of fish is transiting through the sampled location. Predicting anchovy habitats with a classifier, like the XGBoost model or the GAM, can be challenging if the presence of anchovy schools is probabilistic even under suitable environmental conditions. A key problem is that traditional classifiers might struggle to capture the stochastic nature of anchovy appearances, as they typically aim for deterministic outcomes. This means that, even if the environmental conditions are right, the inherent unpredictability of anchovy locations is not easily learned by standard classifiers, which could lead to inaccuracies in habitat prediction. Our clustering method proved to be robust for handling non-deterministic data and, from a statistical perspective, it was better than the other two methods. It also has potential for predicting abundance which is challenging in datasets that are zero-inflated such as for sparsely distributed organisms (Barry and Welsh, 2002; A. Lee-Yaw et al., 2022). However, predicting the abundance of anchovy is beyond the scope of this paper so we only evaluated the clustering probability of occurrence predictions.A caveat of the clustering method is that, even after scaling, it predicted a very narrow range of probabilities, underestimating in regions where anchovy is common and overestimating in areas where anchovy is absent. If a larger dataset is available, this might be improved by defining more water types. Supervised methods such as XGBoost and statistical methods like GAM have been extensively used in habitat modelling already, while unsupervised learning, like clustering, has received less attention. Thus, further development of unsupervised learning methods might be a promising approach for improving habitat models.

## 5 Summary

We simulated anchovy habitat with three models combined with input from a physical-biogeochemical model. We used a method for comparing the three models with observations based on producing a heatmap from the presence-absence data points which allowed us to train the models using brute force grid search. This approach must be combined with expert judgement based on the visualisation of the maps to pick the best maps. The new unsupervised learning method that we implemented was able to produce a map with a similar distribution pattern for anchovies to the more traditional methods based on supervised learning. However, the range of probabilities of occurrence was rather narrow, over- and underestimating the predictions depending on the region. We also produced climate change predictions for anchovies which indicate a reduction in their potential habitat under warming temperatures and weaker winds and a potential increase under environmental conditions with stronger winds. Two of the three habitat models predicted a severe maximum decline by 90% (GAM) and 75% (XGBoost) whereby the clustering model predicted a moderate maximum decline in anchovy occurrence by 20%.

### A Model comparison with observations

In order to compare with observations, phytoplankton concentrations (*P*, in unit: mmol N m^−3^) in the CROCO-BioEBUS model were converted to chlorophyll *a* (*chl*, in unit: mg m^−3^) using a constant chl/N ratio of 1.59 (*chl* = 1.59 *· P*) following Hurtt and Armstrong, 1996 and Gutknecht et al., 2013. Simulated chlorophyll in CROCO-BioEBUS tends to be higher than in observations (Figure 6, note that chlorophyll is shown in log-scale). This discrepancy may arise from two key factors: (1) the constant chl/N ratio used in the model is too high for the Peruvian system; and (2) the absence of iron limitation in the model, which leads to an overestimation of chlorophyll concentrations offshore where, according to observations, phytoplankton growth is limited by iron availability, as discussed in Xue et al. (2022).

**Figure 6.**
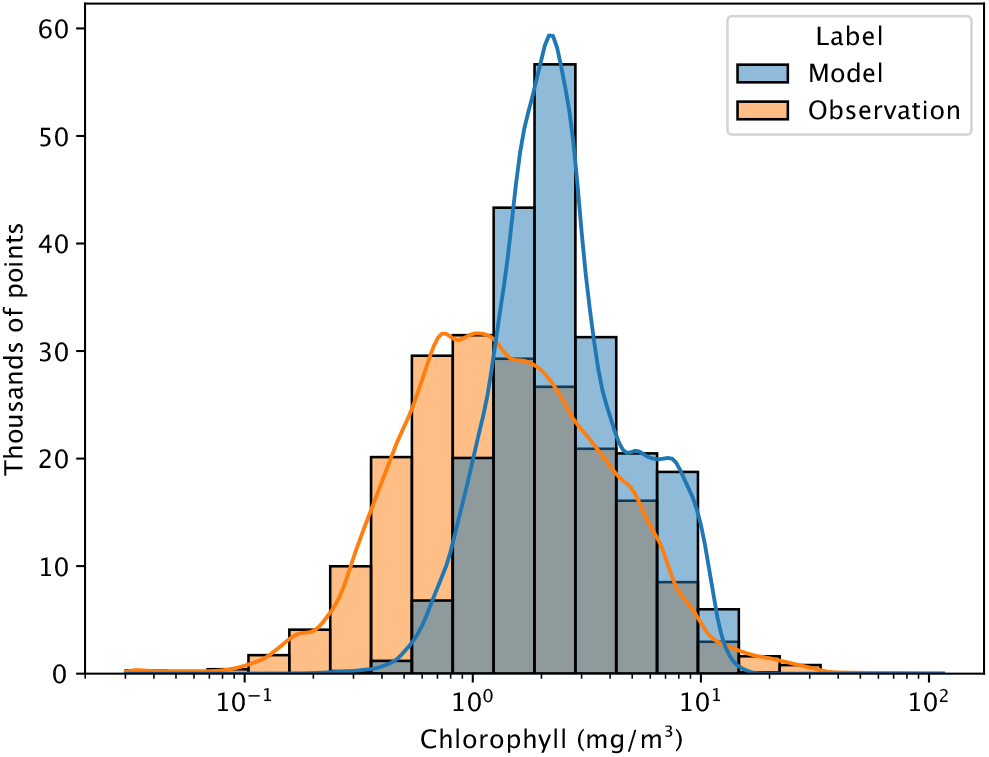
Distribution of chlorophyll *a* in observations and CROCO-BioEBUS output sampled at the same locations.

Fig. 7 shows a higher temperature in observations sampled between January and May than those sampled after May. This is also observed in the CROCO-BioEBUS model although with a less clear differentiation (Figure 8).

**Figure 7.**
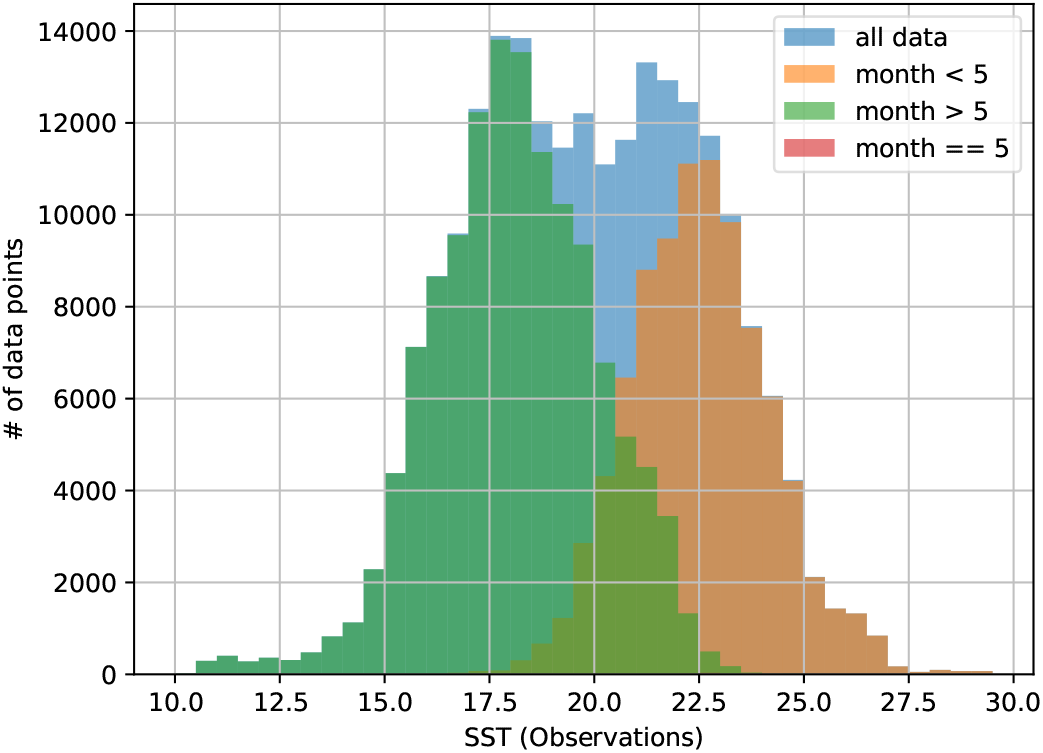
Sea surface temperature distribution in Observations (°C).

**Figure 8.**
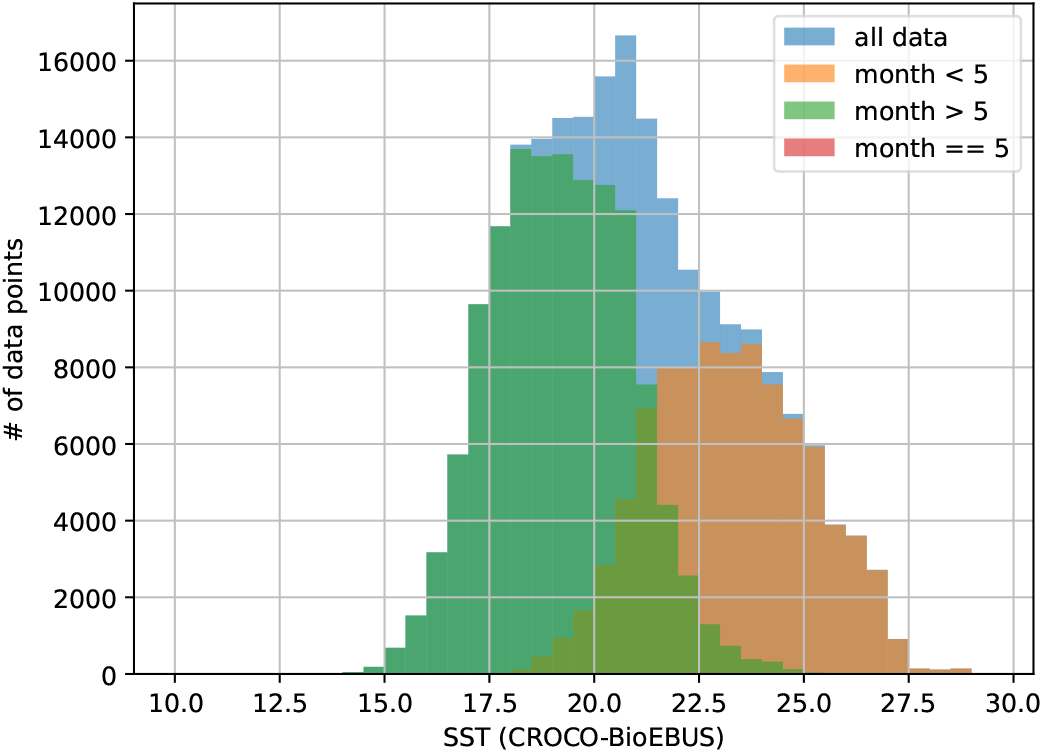
CROCO-BioEBUS sea surface temperature distribution of points sampled at the same location as in observations (°C)

### B XGBoost error type analysis

Out of the environmental variables that we used to predict the habitat of Peruvian anchovies, the depth of the oxycline is the variable that showed the lowest correlation between observations and the physical-biogeochemical model CROCO-BioEBUS (see Section 2.2.1). Therefore, we performed an error type analysis to evaluate whether this variable was responsible of the decrease in the XGBoost test score when running it with predictors from CROCO-BioEBUS with respect to observations (see Section 3.2). Fig. 9 indicates that the model has generally higher accuracy when the oxycline is deep in both observations and the CROCO-BioEBUS model (Figure 9 top-left). This also corresponds to the area with fewer data-points and fewer fish (Figure 9). Since accuracy does not seem to be affected by a higher or lower correlation between observations and CROCO-BioEBUS output (Figure 9 top-left), we cannot conclude that the low correlation between observations and CROCO-BioEBUS has significantly impacted the overall accuracy of the habitat map.

**Figure 9.**
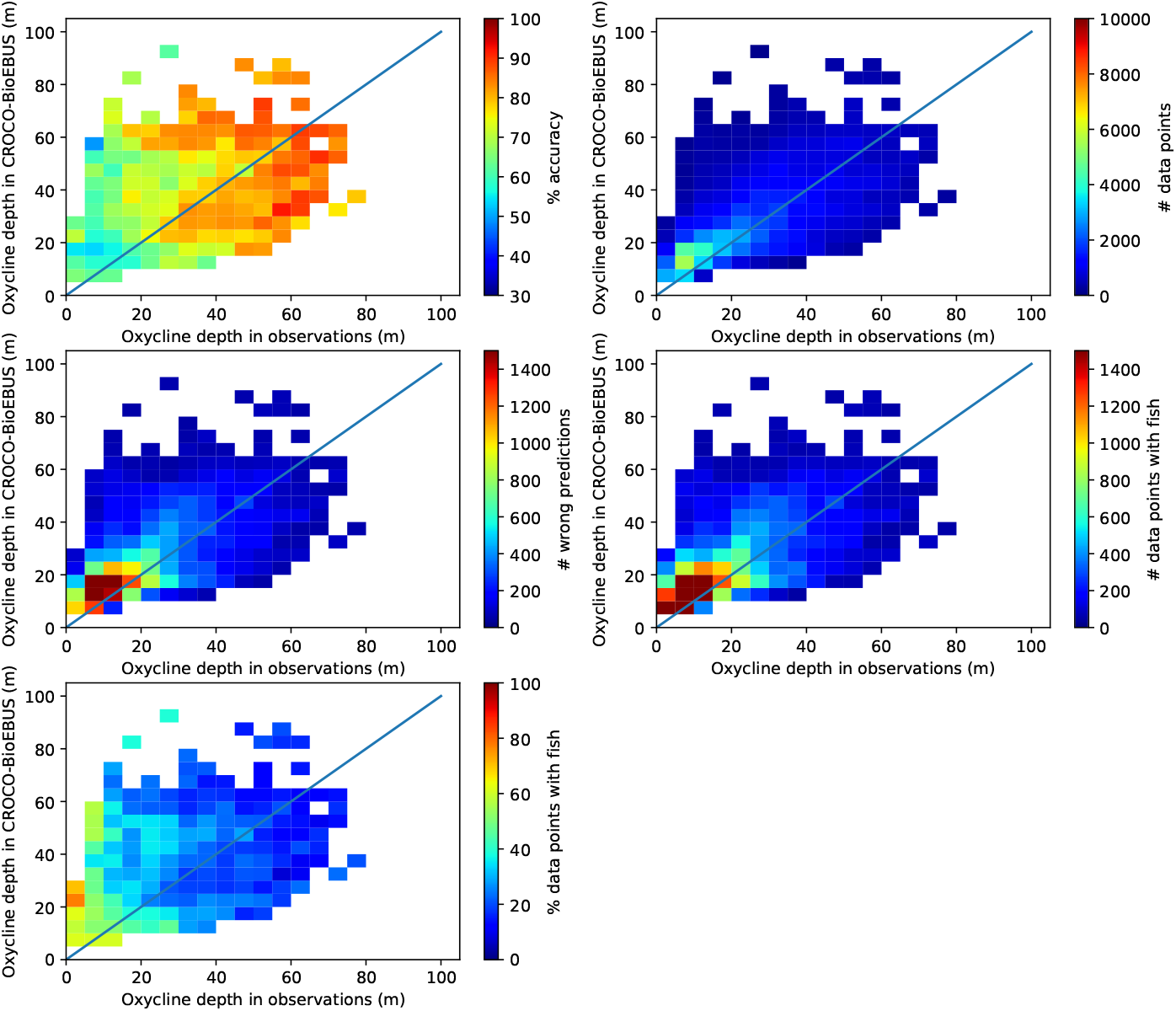
Error type analysis of the depth of the oxycline.

### C Characteristics of the water types

Water types were clustered according to similarities in their environmental characteristics. These are shown in Fig. 10.

**Figure 10.**
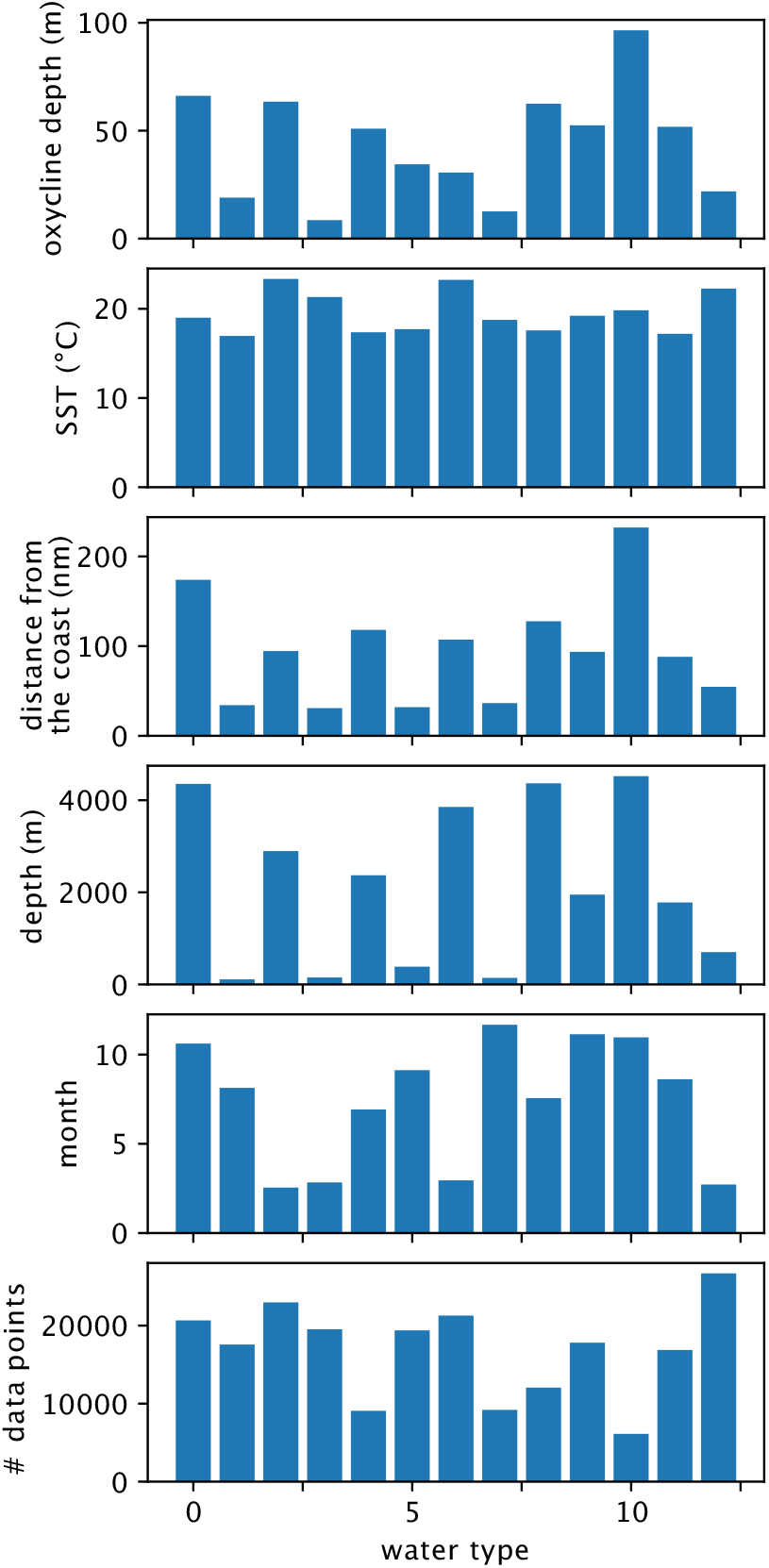
Features of the clustered water types.

## Author contributions

MH: Conceptualising the original idea and designing the study, leading the paper writing, organisation, discussion of results and preparing figures. TX: Conceptualising the original idea and designing the study, running the CROCO-BioEBUS hindcast simulation and preparing the model output, discussion of results, preparing figures and contribution to the manuscript text. JH: Running the CROCO-BioEBUS climate sensitivity simulation, discussion of results, preparing figures and contribution to the manuscript text. MG: Expertise on the northern Humboldt Current System and the Peruvian anchovy, data compilation, discussion of results and contribution to the manuscript text. TK: Setting up and tuning the habitat models, proposing the clustering method and the common method to evaluate all three models, data analysis, discussion of results, preparing figures and contribution to the manuscript text. MH and TK contributed equally to the paper.

## Acknowledgements

The authors thank Daniel Lizarbe and David Moncayo for their valuable feedback and comments to improve this paper. We also thank Anna Akimova and Klaus Huebert for their suggestions. CROCO-BioEBUS simulations were carried out using the computing facilities of the Norddeutscher Verbund zur Förderung des Hoch-und Höchstleistungsrechnens – HLRN. This project received financial support by the Bundesministerin für Bildung und Forschung (BMBF)-funded project GlobalTip: Humboldt-Tipping (01LC2323B).

## Model availability

The CROCO-BioEBUS hindcast output is available at the GEOMAR data management server: https://hdl.handle.net/20.500.12085/b4d40ba5-48ad-48c8-99c4-fc422aa3cebd (Xue et al., 2023). The source code is available and maintained at the official CROCO webiste: https://www.croco-ocean.org/.

